# A BCI-based vibrotactile neurofeedback training improves motor cortical excitability during motor imagery

**DOI:** 10.1101/2021.02.28.433220

**Authors:** N. Grigorev, A. Savosenkov, M. Lukoyanov, A. Udoratina, N. Shusharina, A. Kaplan, A. Hramov, V. Kazantsev, S. Gordleeva

## Abstract

In this study, we address the issue of whether vibrotactile feedback can enhance the motor cortex excitability translated into the plastic changes in local cortical areas during motor imagery (MI) BCI-based training. For this purpose, we focused on two of the most notable neurophysiological effects of MI – the event-related-desynchronization (ERD) level and the increase in cortical excitability assessed with navigated transcranial magnetic stimulation (nTMS). For TMS navigation, we used individual high-resolution 3D brain MRIs. Ten BCI-naive and healthy adults participated in this study. The MI (rest or left/right hand imagery using Graz-BCI paradigm) tasks were performed separately in the presence and absence of feedback. To investigate how much the presence/absence of vibrotactile feedback in MI BCI-based training could contribute to the sensorimotor cortical activations, we compared the MEPs amplitude during MI after training with and without feedback. In addition, the ERD levels during MI BCI-based training were investigated. Our findings provide evidence that applying vibrotactile feedback during MI training leads to (i) an enhancement of the desynchronization level of mu-rhythm EEG patterns over the contralateral motor cortex area corresponding to the MI of the non-dominant hand; (ii) an increase in motor cortical excitability in hand muscle representation corresponding to a muscle engaged by the MI.

## Introduction

Neurofeedback (NF) is a type of biofeedback that employs the registration and real-time feedback of brain activity to users to promote neural self-regulation and improve cognitive control. NF results in neuroplasticity in the trained brain circuit and is being used in novel experimental and clinical applications [1]. Maksimenko et. al. [2, 3] showed that the brain–computer interface (BCI) based biological feedback affects visual perception and prolongs the periods of sustained attention. As a variation of NF, training in motor imagery (MI) based BCI is believed to be a helpful technique in neurorehabilitation therapy of people with impaired motor functions (e.g. patients with tetraplegia, spinal cord injury) and patients with brain injuries (e.g. stroke, amyotrophic lateral sclerosis) [4-6]. Therefore, considerable efforts have been made to develop and study the different paradigms of BCI-based NF for recent reviews, see [6-8].

One of the key components of BCI training is the feedback given to the subject to improve the learning of an MI task and promote motivation and engagement [9]. Traditionally, most MI-based NF systems have used visual feedback due to the simplicity of its implementation and understanding by the user [10-13]. However, in some cases, visual feedback is not suitable, for example, for patients with an impaired visual system [14] or under the circumstances of the visual channel overload [15-18]. For these purposes, other feedback modalities have been explored. For MI-based NF training, closing the sensorimotor loop through haptic feedback can improve motor recovery through enhancing activity-dependent neuroplasticity [19-22]. It has been widely shown that inclusion of anatomically congruent, proprioceptive feedback (such as robotic exoskeleton devices [23-25], functional electrical stimulation [20-22], virtual reality [26-29], vibrotactile stimulation [15,18, 30-33], etc.) induced NF effects superior to non-physiological feedback. Most MI-based NF system use orthosis and exoskeletons as kinesthetic feedback for research [23, 24, 35, 36] or for clinical applications, especially for the rehabilitation of stroke patients [19, 38-41].

Compared to robotic devices, vibrotactile interfaces with proprioceptive feedback functions are more comfortable, affordable, portable, cheaper and easier to implement. However, up until now, vibrotactile feedback has been scarcely explored in BCI-based NF training applications (for a recent review see [42]). The pioneering studies on this type of haptic feedback were conducted by Chatterjee et al. [43] and Cincotti et al. [16]. They showed that users could control MI-BCI using only vibrotactile feedback. Shu et al. [33] proposed the integration scheme of the MI task with the constant tactile stimulation applied to the non-dominant/paretic hand of healthy and stroke individuals. It induced significant enhancement of the contralateral cortical activations during MI of the stimulated hand and, as a result, led to improved of BCI performance. A similar way of applying vibrotactile stimulation for BCI accuracy improvement was presented by Yao et al. [34] for a hybrid BCI system based on a combination of MI with tactile sensation. Together with previous research, their goal was to enhance MI via vibrostimulation [33] and solve the “BCI illiteracy” problem [44]. In another study, Barsotti et al. [32] showed that multisensory (visual and vibrotactile) feedback enhanced the EEG event-related-desynchronization (ERD) during MI and led to higher BCI performance relative to using visual feedback only. In this study, subjects received continuous tendon vibration feedback during MI, according to the online decoded mental state.

It is shown that MI promotes not only ERD but also the enhancement of corticospinal excitability as evaluated from the amplitudes of motor-evoked potentials (MEPs) [45-49] induced by transcranial magnetic stimulation (TMS). TMS is a non-invasive electrophysiological technique that allows studying the excitability of different brain regions [50, 51]. Despite the numerous studies focused on the investigation of the neurophysiological effects induced by the MI-based BCI, little is known about how adding the feedback to BCI training influences cortical excitability. However, it was shown that increased motor cortex excitability during MI BCI-based NF training can promote the plasticity in motor cortical areas, which opens up a wide range of neurorehabilitation therapy technologies.

In our previous research, we proposed the BCI paradigm employing only a vibrotactile channel – without the use of visual control elements (with the eyes closed) [18]. We showed that the characteristics of EEG activity, corticospinal excitability, session-by-session dynamics, and the accuracy of BCI use in this approach were at least no different from those in the classical scheme with visual delivery of stimuli and feedback. In this study, we address the issue of whether vibrotactile feedback can enhance the motor cortex excitability translated into the plastic changes in local cortical areas during MI BCI-based training. For this purpose, we focused on two of the most notable neurophysiological effects of MI – the ERD level and the increase in cortical excitability assessed with navigated transcranial magnetic stimulation (nTMS). The MI (rest or left/right hand imagery using Graz-BCI paradigm) tasks were performed separately in the presence and absence of feedback. To investigate how much the addition of vibrotactile feedback in MI BCI-based training could contribute to the sensorimotor cortical activations, we compared the MEPs amplitude during MI after training with and without feedback. In addition, the ERD levels during MI BCI-based training sessions were investigated.

## Methods

### Subjects

Ten BCI-naive and healthy adults (6 females; age range 18-27 years old, mean ± SD: 22.5 ± 2.3) participated in this study. All subjects were right-handed (mean ± SD: 0.80 ± 0.21 points according to the Edinburgh Handedness Inventory). The protocol was realized in accordance with the Declaration of Helsinki and approved by the Ethical Committee of the Institute of Biology and Biomedicine of the Lobachevsky State University of Nizhny Novgorod. All the participants gave their written informed consent in advance.

### Experimental paradigm

The aim of this study was to apply BCI-based vibrotactile NF training to improve the subject’s motor cortical activations during MI. The experiment consisted of four sessions, each taking place on one experimental day (training session, MI BCI training session without vibrotactile feedback, control session and MI BCI vibrotactile training session). Figure 1 illustrates the experimental paradigm procedure. During each session, subjects were comfortably seated in a reclining chair, with both arms positioned on armrests. The experimental scenarios were displayed on a 27-inch LCD screen located at a distance of 2 meters. Each experimental session lasted 2–2.5 h and was followed by the TMS measurement (except the training session).

**Figure 1.**
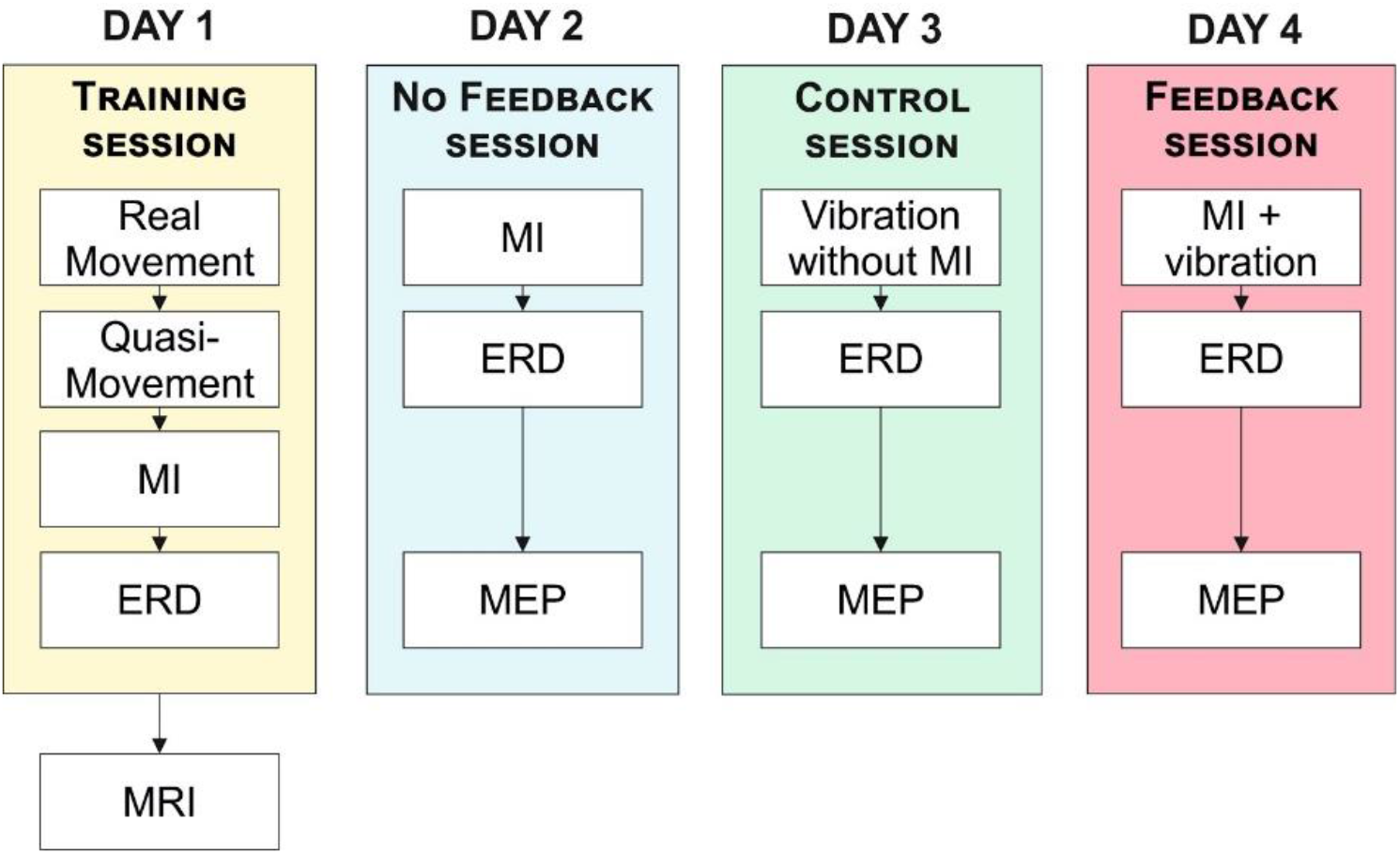
Experimental paradigm. The experiment consisted of four sessions, each taking place on one experimental day. *Day 1* - during the first training session, the subjects trained kinesthetic left-or right-hand MI. Trained subjects underwent magnetic resonance imaging (MRI) of the head. *Day 2* - the second session featured the BCI-based training without vibrotactile feedback utilizing the classical Graz MI-BCI protocol. *Day 3* - the third session was the control session. The subjects were asked to perform a rest command regardless of the cue presented. *Day 4* - the fourth session featured vibrotactile NF training. During on- line testing, if the MI or rest task was correctly classified, vibrotactile feedback was applied to subjects. MEPs measurement was performed 30 min after the last BCI training run on *Days 2, 3, 4*.

In the first training session, the subjects trained in kinesthetic left-or right-hand MI. The training consisted of three sequential steps: hand movements, quasi-movements (muscle tension is not observed visually), and MI. The subjects were asked to clench their hands into a fist. The success of training was determined by the ERD power in the sensorimotor cortex during MI. The training progress was represented to the subjects by voice. Trained subjects underwent magnetic resonance imaging (MRI) of the head to build a 3D model of the brain structures for nTMS.

The second session was the BCI-based training without vibro feedback utilizing the classical Graz MI-BCI protocol [52]. This session consisted of three 5 min 5 sec test runs of 30 trials each. Figure 2a illustrates the MI-BCI protocol. Subjects performed one of three commands: kinesthetic MI of left or right hand (fist clenching) or rest when the subject had to concentrate on their breathing. Subjects were asked to execute the command after they would see a visual cue (Fig. 2a). Each command was repeated 10 times during a testing run with randomized order. The duration of each command was 5 s, and the interstimulus interval was 5 s (Fig. 2a). During the interstimulus intervals, the subjects were allowed to blink and swallow. Between testing runs, subjects could rest as long as they felt necessary. The classifier analyzed EEG every 500 ms. MEPs measurement was performed 30 min after the last BCI training run.

**Figure 2.**
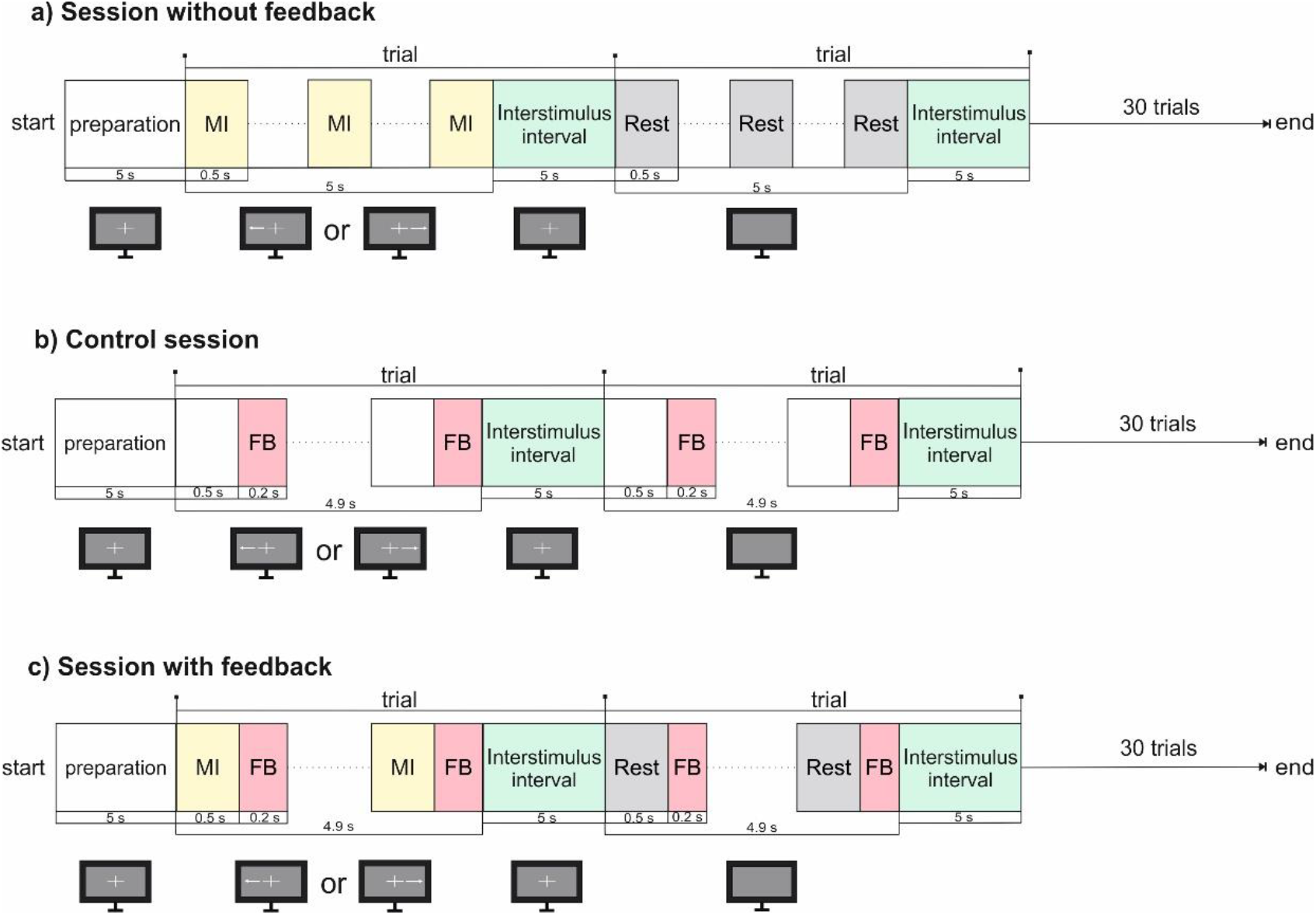
Timing of MI-BCI sessions without (a) and with (c) vibrotactile NF. (b) Timing for control session. MI - motor imagery, FB - vibrotactile feedback.

The third session was the control. The subjects were asked to perform rest command regardless of the cue presented. During each command, the subjects were stimulated via vibrotactile actuators for 200 ms with the interstimulus interval of 500 ms (Fig.2b). The control session consisted of three runs of 30 trials each and was followed by nTMS measurements.

Figure 2c illustrates the experimental protocol for the fourth vibrotactile NF training session. The structure of the protocol was the same as in the second session of MI BCI training. There were 3 test runs in this session. Each run consisted of 30 command trials (kinesthetic MI of left or right hand or rest) with randomized order. As the visual cue appeared, the subjects performed the mental task for 4.9 sec. During the mental task, the online performance was tested by classification of EEG in time windows of 500 ms. During online testing, if the MI or rest task was correctly classified, vibrotactile feedback was applied for 200 ms immediately after the classification period ended. MEPs measurement was performed 30 min after the last vibrotactile NF training run.

### EEG recording and classification

For data acquisition, we used a 48-channel NVX-52 amplifier (MKS, Zelenograd, Russia). The EEG data were recorded from 6 standard Ag/AgCl electrodes (C5, C3, C1, C2, C4, C6), placed according to the international 10-10 system (Fig. 3a). The earlobe electrodes were used as a reference. The grounding electrode was placed on the forehead. All electrode impedances were kept below 15 kΩ. EEG was digitized with a signal sampling frequency of 1 kHz and filtered in the frequency range from 1-30 Hz with a 50 Hz Notch filter.

**Figure 3.**
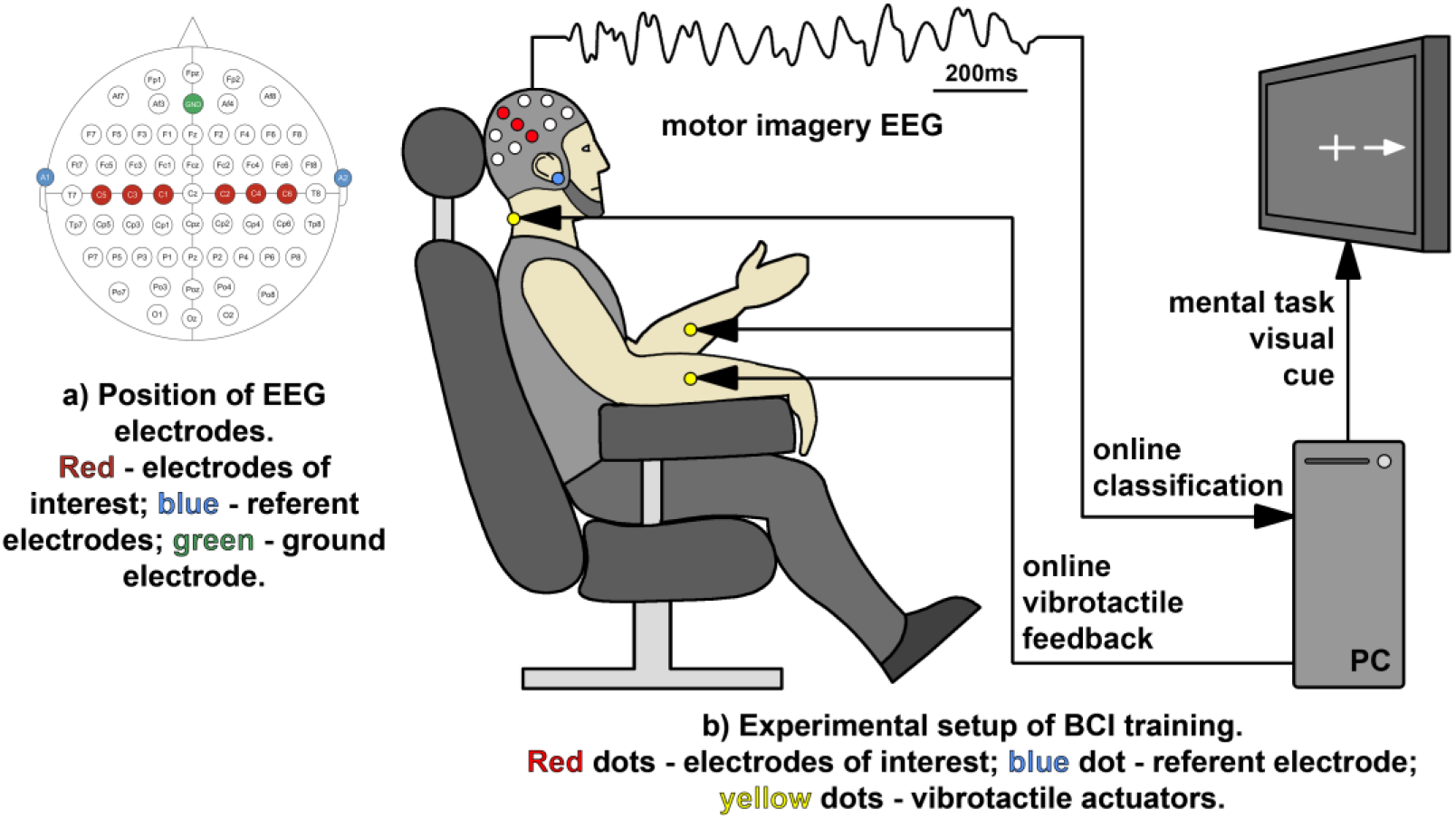
Experimental setup for MI-based BCI vibrotactile NF training session. (a) EEG electrode distribution in the 10–10 system. (b) Subjects were comfortably seated in a reclining chair, with both arms positioned on armrests. The vibrotactile actuators were attached to the right and left forearm and to the back of the neck in order to provide real-time feedback of the successful decoding of MI of the right and left hand and of the rest task, respectively.

The EEG data of the first five seconds of each trial were extracted for pattern classification. The raw signals were bandpass filtered using the 4th-order Butterworth filter at 7-16 Hz, followed by the calculation of the coefficients for the common spatial pattern (CSP) filter [53]. We used the linear discriminant analysis (LDA) method to classify the EEG patterns [54]. During online classification, the classifier analyzed EEG every 500 ms. In the sessions with vibrotactile stimulation, the time windows of feedback application were removed from the classification. The second and fourth experimental sessions started with the training of the LDA classifier. For classifier training, we used one run, the timing of which repeated the test run in the BCI-based training without feedback session. The custom Python script was used for data recording, classification and feedback control.

The average classification accuracy for three classes (the MI of left/right hands and the rest task) was calculated as a percentage of correct classification time (7 times in one trial).

To estimate ERD levels during MI, signals corresponding to the rest task were taken as the reference state. The raw EEG data were spatially filtered using the Surface Laplacian [55] for all the channels. Then, the power spectral density was constructed for each channel with a step of 1 Hz, and ERD was calculated as the difference between the signal powers during MI and the rest signal, which was divided by the signal power corresponding to the rest task. Then for each subject, an individual frequency range was chosen in which the peak ERD was most often encountered. For statistical analysis, in this individual frequency range, one value of the maximum ERD was chosen in each 500 ms MI time window for each electrode. In the sessions with feedback, the time intervals of vibro stimulation were removed from the analysis.

### Vibrotactile Feedback

A flat linear resonant actuator (3 V, diameter 10 mm) was used for tactile stimulation. The actuator was positioned on the right and left forearm and on the back of the neck to feedback the successful decoding of MI of the right and left hand and the rest task, respectively (Fig. 3b). Actuators were fixed on the skin using Velcro tapes. The actuators with selected vibration parameters were operated by the Arduino Nano microcontroller connected to a PC via COM-port. To confirm a correctly classified command, a vibration signal lasting 200 ms was applied to a subject. The pattern and intensity of vibration were chosen to give the user the most pleasant and distinct feedback. A vibrotactile stimulation with the selected amplitude and duration was tested on subjects before the experiment to ensure the stimulation can be perceived, is comfortable, and does not prevent them from freely performing the required MI tasks.

### MEPs Measurement

After control and BCI-based training sessions, if needed, EEG caps were removed from the subjects, and they were prepared for MEPs measurements. The preparation took about 30 min. Cortical excitability was only measured for the right hand, as a measurement for both hands would have made the experiment too long for the participants. MEPs were obtained using nTMS. For navigation, we used the Localite TMS Navigator system (Localite, Germany) with individual high-resolution 3D brain MRI. For matching each subject’s head and MRI data, we used three landmarks: left and right lateral canthi and nose bridge. The navigator oriented individual 3D-MRI data to the subject’s head through infrared tracking using a marker with spheres coated with a reflective surface.

While applying nTMS, muscle activity was continuously monitored using electromyography (EMG). EMG was recorded with pair of Ag/AgCl hat-shaped electrodes (COVIDIEN, USA) from flexor digitorum superficialis (DS) muscle on the right hand. A ground electrode was placed on the left forearm. The skin under the electrodes was prepared with alcohol cotton swabs. Electrode impedances were kept below 15 kΩ. Signal was digitalized at 500 Hz using the Neuron-Spectrum-5 amplifier (Neurosoft, Ivanovo, Russia) and filtered with a 50 Hz Notch filter.

Single-pulse TMS was applied through a figure-of-eight shaped coil (7 cm diameter) connected to a Neuro MS/D magnetic stimulator (Neurosoft, Ivanovo, Russia) to the motor cortex, in the optimal location to measured MEPs in the contralateral DS of the right hand (“hotspot”, [56]). DS hotspot for each subject determined during the second session was recorded by the navigator and used in subsequent experimental days. The coil was positioned tangentially to the skull with the handle pointing back and away from the midline by 45°. The resting motor threshold (RMT) was determined as the minimum TMS intensity able to evoke MEPs larger than 50 μV in at least five out of ten consecutive trials [56]. TMS intensity was fixed at 110 % of RMT (37 ± 5%) and kept constant. Figure 4 illustrates the experimental setup.

**Figure 4.**
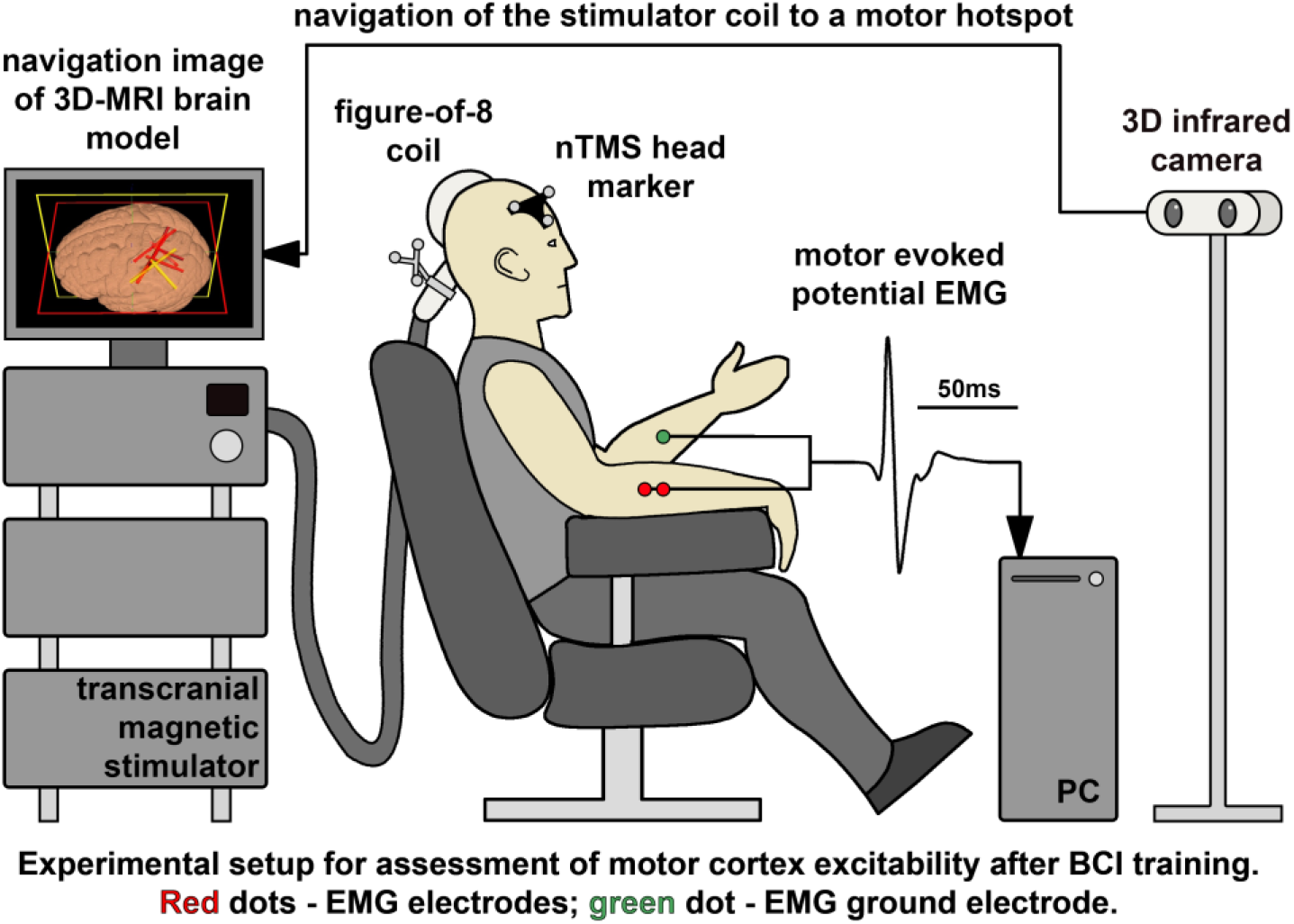
Experimental setup for MEPs measurement after BCI NF training. To obtain MEPs, nTMS oriented by individual 3D-MRI data were used. During rest and motor imagery TMS was applied to the sensorimotor cortex of the right hand. MEPs were recorded by EMG data from the flexor digitorum superficialis muscle on the right hand.

First, we recorded MEPs during the rest condition, when subjects were comfortably seated in a reclining chair, with both arms positioned on armrests, eyes open, and fully relaxed. Then, subjects were asked to kinesthetically imagine repetitive right fist clenching. We asked the subjects to try to synchronize the moments of the fist clenching imagery with the TMS stimulus. There were three runs at rest and during the MI task. Each run was 2 min long and consisted of 60 stimuli with 0.5 Hz frequency. There was a break of 1 min between runs. EMG data were monitored in real-time during the entire duration of the MEPs measurement to ensure that the coil orientation was stable across stimulus delivery, and was recorded for offline analysis. We used Neuron-Spectrum.NET software to measure the amplitude of MEPs from peak-to-peak. MEPs during MI were normalized to median MEPs amplitude at rest on the same session.

### Statistical analysis

The significance of differences between sessions was assessed using pairwise comparison with Wilcoxon signed-rank tests. Statistical analysis of the individual subject’s data was performed using a pairwise Mann-Whitney U-test. The false discovery rate method was used for multiple testing adjustments. Differences were determined to be statistically significant at p < 0.05. All analyses were made in R 4.0.2 software (R Core Team, 2020). Descriptive statistics of the ERD levels and MEPs amplitudes per group are represented as “M [Q1; Q3]”, where M—median, Q1—first quartile (quantile 0.25), and Q3—third quartile (quantile 0.75) of the group samples.

## Results

To study how much the vibrotactile NF training could contribute to the sensorimotor cortical activations during MI, we employed three characteristics: BCI performance as classification accuracy, the ERD level, and the MEPs amplitudes.

### Classification accuracy

Classification accuracy of three mental tasks (left, right hand MI and rest) during BCI training was compared for two variants: with and without vibrotactile feedback. Individual average accuracies for each subject are depicted in Fig. 5. All subjects showed classification accuracy above the chance level of 0.33 for three classes. Vibrotactile feedback applied to the subjects did not lead to statistically significant accuracy changes (*p=*1). For BCI without feedback, the mean accuracy was 61%, SEM=2.51% (n=10), ranging from 51.7% to 76.2%; and with feedback, the mean accuracy was 61.5%, SEM=2.75% (n=10), ranging from 45.1% to 72.6%. Accuracy was calculated for 500 ms intervals of EEG. In the session with feedback, the time windows of vibro stimulation presence were removed from the classification.

**Figure 5.**
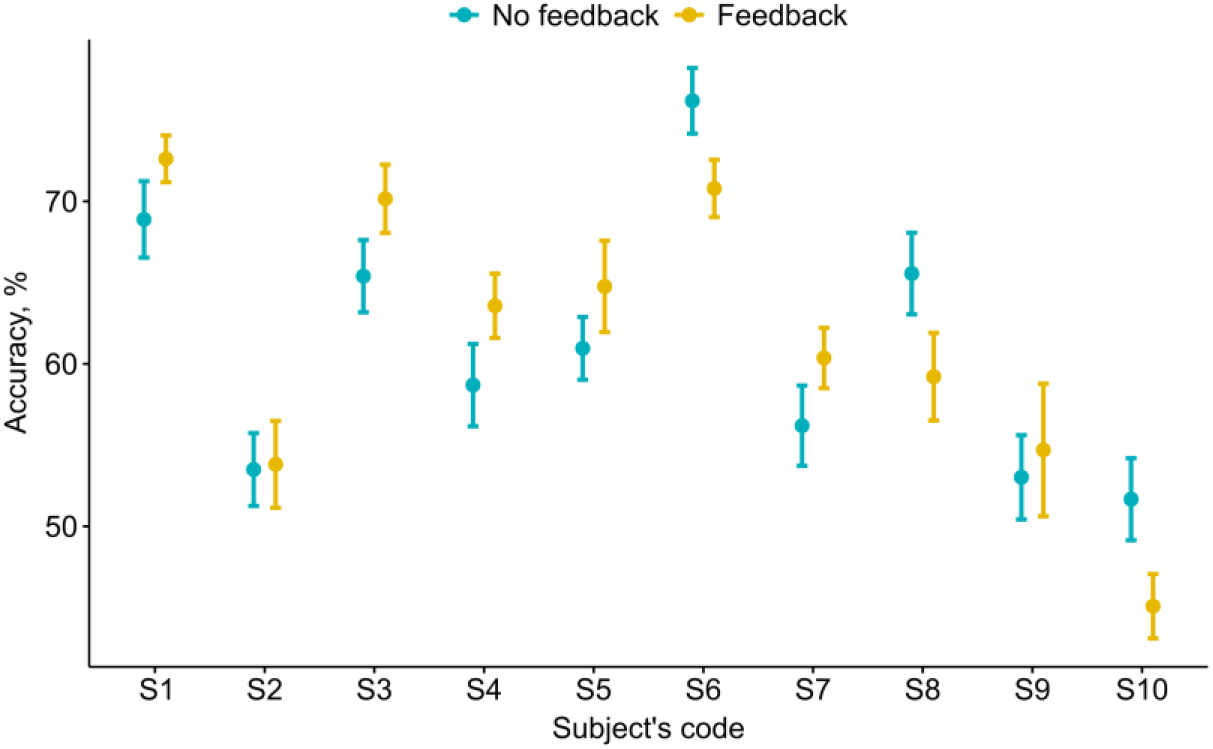
Classification accuracy MI-based BCI for three classes (left/right hand MI and rest) with and without vibrotactile feedback. Mean values are shown with the standard error of the mean indicated by whiskers (n=90).

### ERD levels analysis

ERD levels were estimated for all the subjects using the same EEG samples as for the classification accuracy calculation. The analysis also included the control session data. EEG analysis in the control session showed that vibrotactile stimulation did not induce pronounced ERD over the sensorimotor cortex. Figure 6 depicts the peak ERD levels over the electrodes C3 and C4 for MI training without/with feedback and for the control session. The maximum ERD levels in the control session were -59.2[-65.6; -52.3]% and -62.7[-78.2; -53.9]% for the C3 and C4 electrodes, respectively, and statistically differed from the ERD induced by the MI (*p<*0.01). For the peak ERD levels over the contralateral electrode C3 during right hand MI, no statistically significant differences were found for BCI training without (−82.1[-86.5; -77.0]%) as compared to with (−86[-90.9; -75.9]%) feedback (*p=*0.193). However, applying the vibrotactile feedback during left hand MI training in BCI evoked a significant decrease of the ERD over the electrode C4 (*p=*0.027). The peak ERD level on C4 was -84.5[-90.1; -80.9]% and -89.7[-91.5; -85.7]% during left hand MI training without and with feedback, respectively.

**Figure 6.**
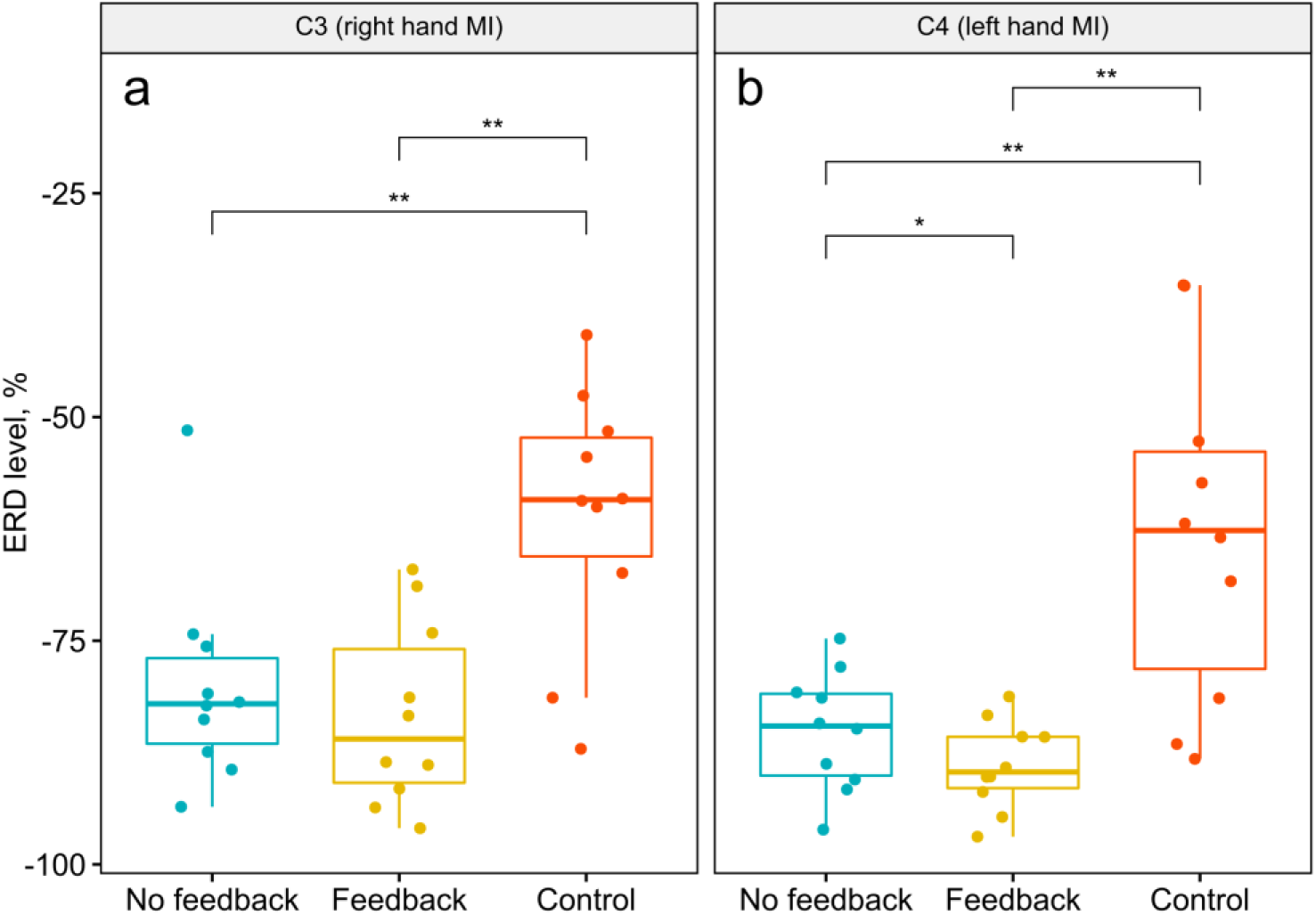
The peak ERD levels over the contralateral C3 and C4 electrodes for MI training with and without feedback and for the control session for all subjects (n=10). The horizontal red lines indicate the median, first and third quartile values, while the whiskers indicate the 1.5 IQR, each dot represents one subject. * -*p<*0.05; ** -*p<*0.01.

Figure 7 illustrates the ERD levels during MI over the electrodes C3 and C4 of all subjects for BCI-based training sessions without and with applying vibro feedback. Three subjects (S4, S6, S8) showed a statistically significant decrease in the ERD level for both right-and left-hand MI training sessions in the presence of feedback. For subjects S1, S2, and S5, feedback led to a significant decrease of the ERD during only one hand MI (for S1, S2 -right hand MI; for S5 -left hand MI). Subjects S9 and S10 did not show significant changes in the ERD depending on the presence of feedback. The feedback evoked a significant increase in the ERD level for S3 during left hand MI and for S7 during the right-hand MI.

**Figure 7.**
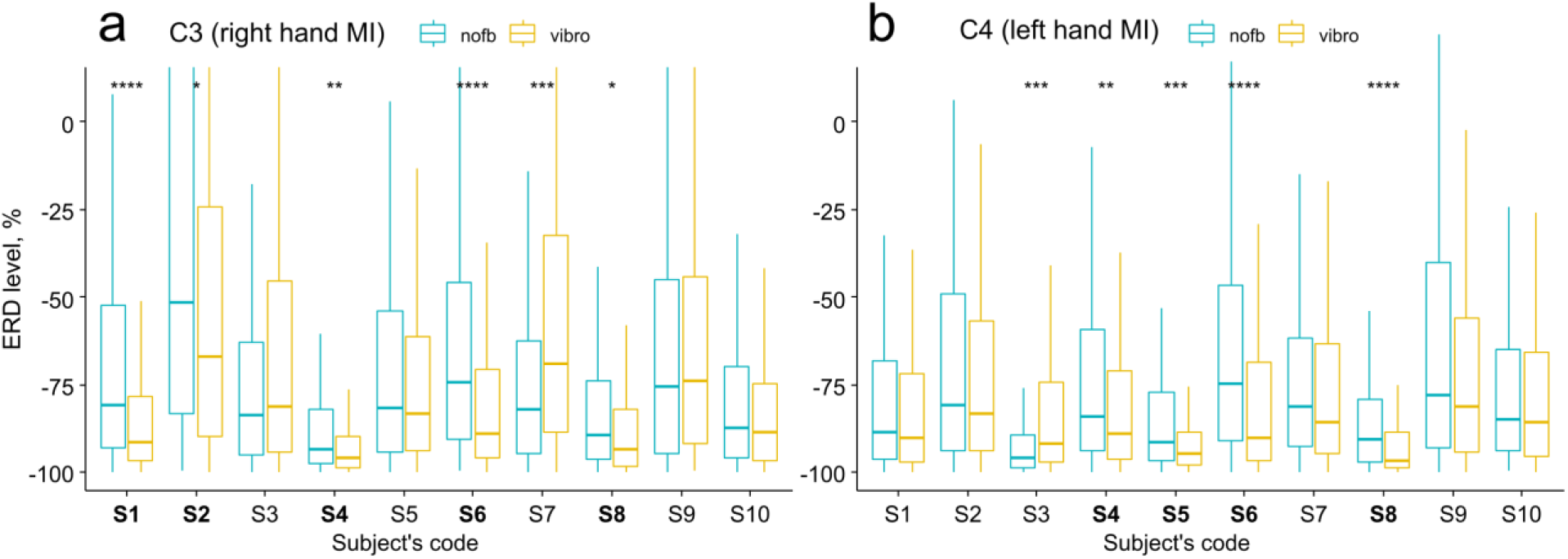
Comparisons of the peak ERD levels during right hand MI training over the contralateral C3 electrode and during left hand MI training over the electrode C4 without and with applying vibrotactile feedback across all subjects (n=210). Subjects with a significant increase in the ERD in the presence of feedback are marked in bold. * -*p<*0.05; ** -*p<*0.01; *** -*p<*0.001; **** -*p<*0.0001.

### MEPs Measurement

Using nTMS, cortex excitability measurement was performed 30 min after all BCI-based training sessions (excluding the first one). MI of right fist clenching resulted in a significant (*p<*0.001) MEPs amplitude increase (Fig.8) in flexor DS muscle on the right hand for all subjects in all sessions with median values ranged from 216% to 311% of the referential state.

**Figure 8.**
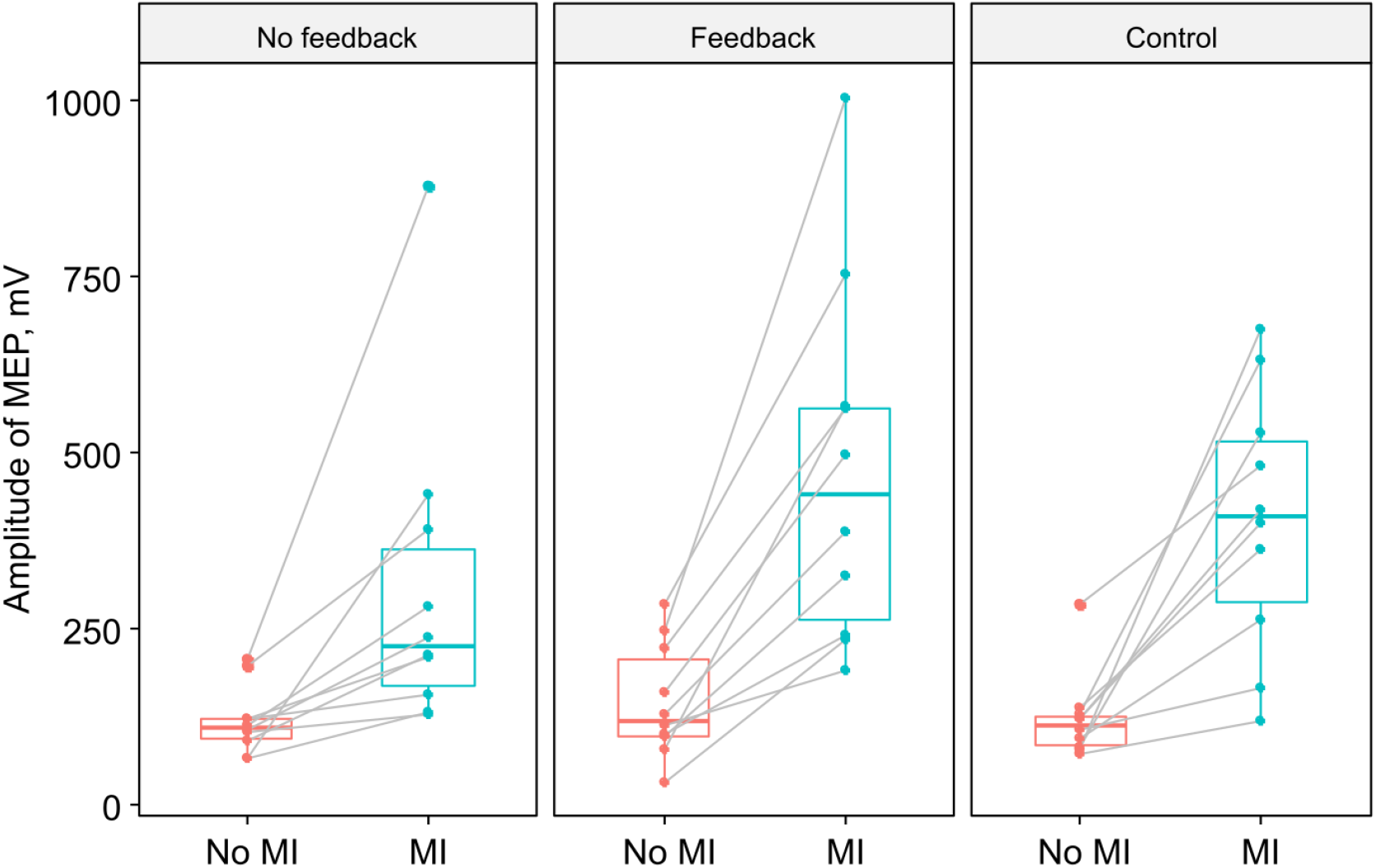
Increase of MEPs amplitudes in flexor digitorum superficialis (DS) muscle on the right hand during MI for all subjects (n=10) after BCI-based training sessions. The lines show individual changes for the subjects.

Figure 9 illustrates the results of within-subjects comparison analysis of MEPs increases during MI after BCI-based training without/with feedback and in the control. Feedback presence in the MI BCI-based training led to statistically significant MEPs amplitude increase for seven subjects (S1, S2, S4, S5, S8, S9, S10) compared to training without feedback. Five subjects (S3, S6, S7, S9, S10) showed the maximum of MEPs increase after the control session. The correlation analysis between the MEPs amplitude increase and the ERD level did not reveal a statistically significant relationship in the BCI-based training sessions either without (Pearson’s r = -0.16, *p=*0.66) or with (Pearson’s r = -0.2, *p=*0.58) feedback.

**Figure 9.**
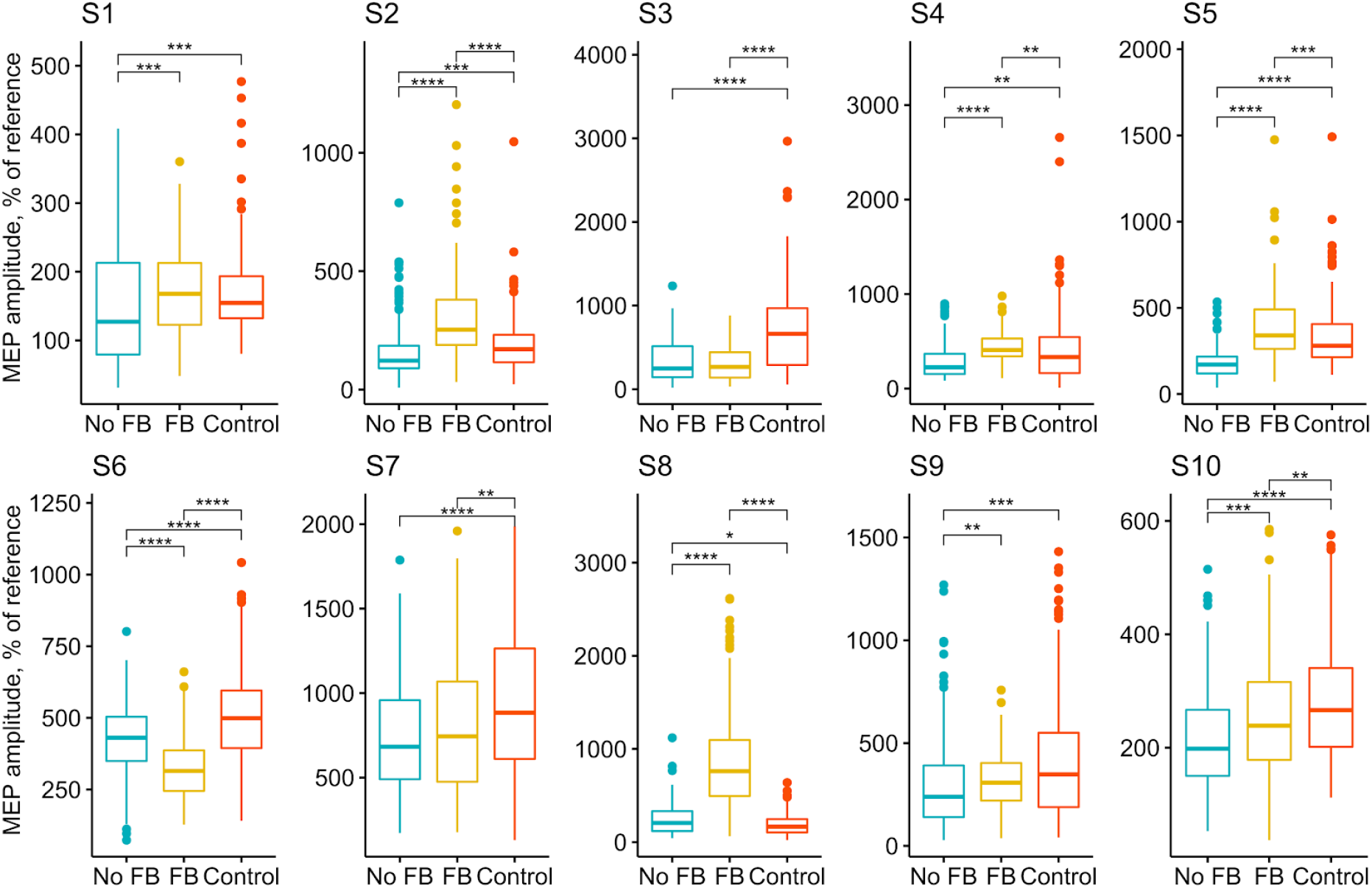
Comparison of MEPs amplitude increase in DS during MI across all subjects after three different types of BCI-based training: without/with feedback and in the control (n=183). * -*p<*0.05; ** -*p<*0.01; *** -*p<*0.001; **** -*p<*0.0001.

For all subjects, MI after BCI-training without feedback resulted in a minimum MEPs increase (216[178; 246]%) compared to the session with vibrotactile feedback and control (*p<*0.05) (Fig. 10). Analysis for the subject group did not reveal a significant difference between amplitude MEPs increase during MI after BCI-training with feedback (311[257, 391]%) and control session (307[195; 461]%) (*p=*0.922).

**Figure 10.**
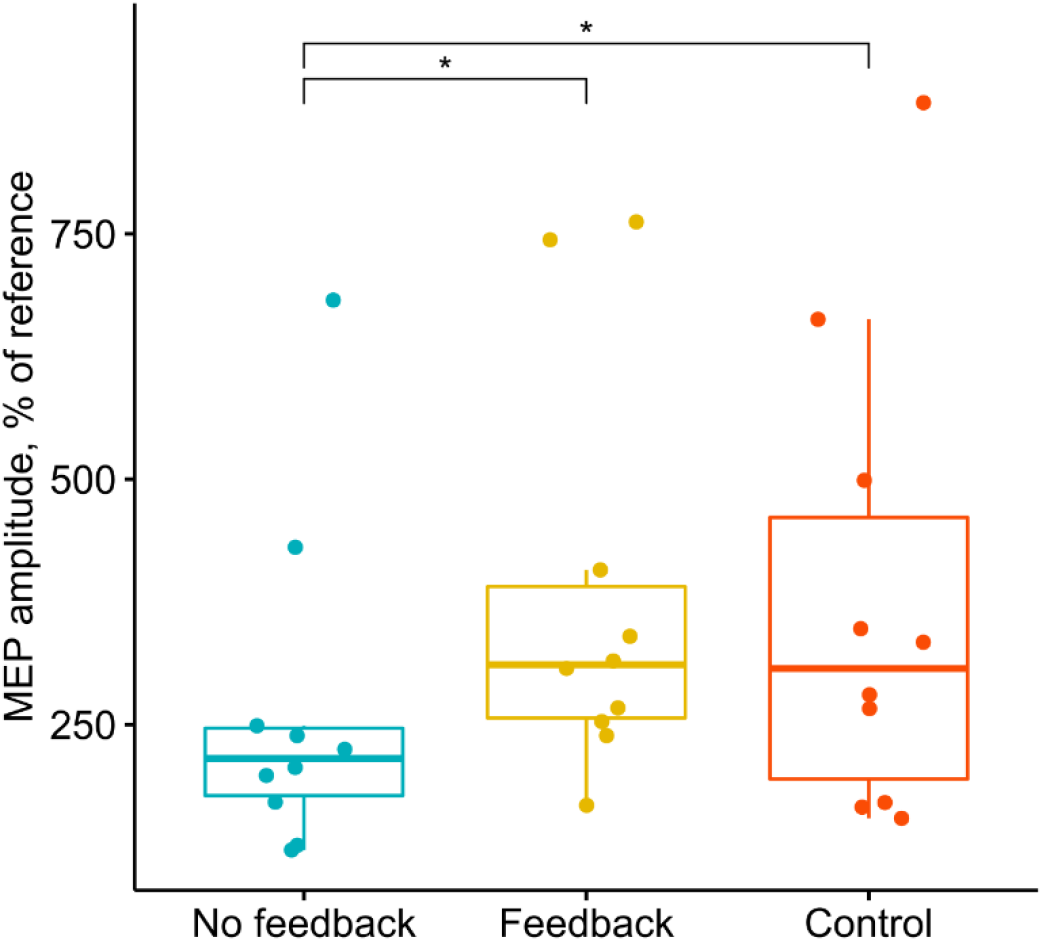
Comparison of amplitude MEPs increase in DS during MI for all subjects after three different types of BCI-based training: without/with feedback and in the control (n=10). * -*p<*0.05.

## Discussion

In this study, we used nTMS and the ERD levels analysis to investigate the impact of introducing vibrotactile neurofeedback to the MI-based BCI training on motor cortical activations in healthy volunteers. Our findings provide evidence that applying tactile feedback during MI leads to (i) an enhancement of the desynchronization level of mu-rhythm EEG patterns over the contralateral motor cortex area corresponding to the MI of the non-dominant hand; (ii) an increase in motor cortical excitability in hand muscle representation corresponding to a muscle engaged by the MI.

### BCI performance

In our study, the average online classification accuracy was 61%, which is significantly higher than chance level 33% for the three-state selection paradigm. This value is not high, nevertheless, it is consistent with the previous studies [19, 31, 32, 57]. The relatively low correct rate achieved could be explained by the fact that the online classification of MI whithin short time windows (500 ms in our study) is more challenging compared with the full several-seconds-long trial classification commonly used in similar studies. When vibrotactile feedback was provided during each trial and not at the end of each trial as previously suggested [58], and was synchronized with the subject’s MI, the resulting haptic stimulation turned out to be more congruent and resembling natural bio-feedback.

The application of vibrotactile feedback did not influence the classification accuracy, which is consistent with our previous studies [58] and with other studies, which showed that the presence of the feedback does not always lead to an improvement of recognition accuracy [25, 59]. However, there are several studies on using vibrotactile stimulation in a different paradigm that were able to provide evidence that MI-based BCI performance can be enhanced via tactile input [32-34].

The proprioceptive input, e.g., vibrotactile stimulation, itself can evoke somatosensory event-related potentials and modulate the cortical activity in a similar way to MI, but independently of any volitional subject intention [60-63]. Usually, the effect of proprioceptive input on brain activity is investigated during the period of haptic feedback application [19, 24, 32, 34], which potentially leads to overlap of the MI-related cortical activity by the additional input of the feedback modality. In our study, we tried to avoid it by taking into account the task-related cortical activity in the absence of feedback only.

### ERD levels analysis

Vibrotactile feedback during MI training induced significant enhancement of ERD activity only for non-dominant, left hand over contralateral motor cortex area measured in C4 electrode. Although the ERD reaction corresponding to the right-hand MI over contralateral C3 also became stronger, it was not statistically significant due to a large variation between subjects. A more noticeable feedback impact on the ERD during left hand MI could be explained by handedness, which resulted in asymmetrical cortical activations between dominant and non-dominant hand MI [64]. The asymmetrical cortical activations between different hands MI may reduce the ERD lateralization. A previous study showed that non-dominant hand MI evoked stronger ERD in the ipsilateral sensorimotor cortex than in the contralateral sensorimotor cortex [65]. Mizuguchi et al. [66, 67] and Shu et al. [33] revealed that applying the tactile stimulation to the imagined hand can enhance the contralateral cortical activations. Thus, the vibrotactile feedback impact on the ERD during non-dominant hand MI could be more expressed.

Feedback-induced increase in the ERD levels did not lead to improved MI BCI classification accuracy. This fact could be explained as the nonlinear relationship of these two characteristics since feature extraction for the MI classification using CSP takes into account the pattern of EEG activity at all electrodes, not just at C3 and C4.

### Cortex Excitability Measurement

In line with previous studies [45, 46, 68, 69], we saw that MEPs were significantly enhanced during the hand MI task in all subjects. In this study, we found clear evidence that including the vibrotactile feedback into MI training significantly increased the motor cortical excitability. For seven subjects out of 10, MI practice with feedback resulted in a significant MEPs increase compared to MI after training without feedback. For two subjects (S3, S7) feedback application did not lead to statistically significant MEPs changes. For the subject S6, MI training with feedback led to the MEPs decrease.

Interestingly, five subjects (S3, S6, S7, S9, S10) showed the maximum MEPs facilitation after the control session. This is probably due to the fact that in the control session vibro stimulation was applied constantly with an interval of 500 ms (simulating 100% accuracy of the MI task recognition). This led to the fact that the number of tactile stimuli in the control session was about 40% higher than in the MI BCI training session. Such intense proprioceptive input can evoke strong activation of cortical areas.

We did not find the correlation between the MEPs amplitude increase and the ERD levels. Although subjects (S1, S2, S4, S8) with the strongest ERD level tended to have a higher increase in cortical excitability, a statistically significant correlation was not observed. This finding is consistent with the similar study of Kaplan et al. [49]. This can be explained by the fact that the MI-induced ERD is an indicator of the general inhibitory input into the vast cortical areas. In contrast, motor cortical excitability represents the state of the particular neuronal circuit corresponding to a discrete muscle. It was believed that MI promotes activation of local cortical pathways engaged in the imagined movement, but does not necessarily affect the general inhibitory output of thalamocortical circuits.

## Conclusion

In this work, we explored the effects of vibrotactile feedback on MI BCI-based training. We found that the addition of the vibrotactile feedback in the MI training paradigm significantly enhances the motor cortical excitability and the contralateral ERD level for non-dominant hand MI. Thus, integrating MI training with vibrotactile feedback can lead to plastic changes in the motor cortex. Finally, our results demonstrate the benefits of using the BCI-based vibrotactile neurofeedback training for recovery of motor function, e.g., after stroke. The findings of our work could be relevant for improving training protocols for rehabilitative interventions after damage to motor cortical areas.

